# Delta and theta neural entrainment during phonological and semantic processing in speech perception

**DOI:** 10.1101/556837

**Authors:** Guangting Mai, William S-Y. Wang

## Abstract

Neural entrainment of acoustic envelopes is important for speech intelligibility in spoken language processing. However, it is unclear how it contributes to processing at different linguistic hierarchical levels. The present EEG study investigated this issue when participants responded to stimuli that dissociated phonological and semantic processing (real-word, pseudo-word and backward utterances). Multivariate Temporal Response Function (mTRF) model was adopted to map speech envelopes from multiple spectral bands onto EEG signals, providing a direct approach to measure neural entrainment. We tested the hypothesis that entrainment at delta (supra-syllabic) and theta (syllabic and sub-syllabic) bands take distinct roles at different hierarchical levels. Results showed that both types of entrainment involve speech-specific processing, but their underlying mechanisms were different. Theta-band entrainment was modulated by phonological but not semantic contents, reflecting the possible mechanism of tracking syllabic- and sub-syllabic patterns during phonological processing. Delta-band entrainment, on the other hand, was modulated by semantic information, indexing more attention-demanding, effortful phonological encoding when higher-level (semantic) information is deficient. Interestingly, we further demonstrated that the statistical capacity of mTRFs at the delta band and theta band to classify utterances is affected by their semantic (real-word vs. pseudo-word) and phonological (real-word and pseudo-word vs. backward) contents, respectively. Moreover, analyses on the response weighting of mTRFs showed that delta-band entrainment sustained across neural processing stages up to higher-order timescales (~ 300 ms), while theta-band entrainment occurred mainly at early, perceptual processing stages (< 160 ms). This indicates that, compared to theta-band entrainment, delta-band entrainment may reflect increased involvement of higher-order cognitive functions during interactions between phonological and semantic processing. As such, we conclude that neural entrainment is not only associated with speech intelligibility, but also with the hierarchy of linguistic (phonological and semantic) content. The present study thus provide a new insight into cognitive mechanisms of neural entrainment for spoken language processing.

**Highlights:**

- Low-frequency neural entrainment was examined via mTRF models in EEG during phonological and semantic processing.
- Delta entrainment take roles in effortful listening for phonological recognition
- Theta entrainment take roles in tracking syllabic and subsyllabic patterns for phonological processing
- Delta and theta entrainment sustain at different timescales of neural processing

## 1. Introduction

Research into how speech acoustic properties are processed by the human brain is key to understanding neural mechanisms of speech and language perception. An important topic that recent research has focused on is to examine how speech temporal modulations are tracked and encoded through brain oscillatory activity (i.e., neural entrainment; see reviews: Giraud and Poeppel, 2012; Ding and Simon, 2014). This is because low-frequency envelope modulations (typically < 10 Hz) are critical acoustic contributors to human speech recognition (Drullman et al., 1994; Shannon et al., 1995; Arai et al., 1999; Swaminathan and Heinz, 2012). Neural entrainment of low-frequency envelopes has been suggested to serve as one of the neural mechanisms of sustaining speech comprehension (Ahissar et al., 2001; Ding and Simon, 2014).

Recent neurophysiological studies using magnetoencephalography (MEG) and electroencephalography (EEG) have shown that entrainment of low-frequency neural oscillations to speech envelopes at the corresponding modulation rates is associated with speech intelligibility (MEG: Peelle et al., 2013; Doelling et al., 2014; EEG: Vanthornhout et al., 2018). Specifically, Peelle et al. (2013) manipulated speech intelligibility by changing the spectral resolution (i.e., number of frequency bands) of noise-vocoded sentences. They found that phase coherence between MEG and acoustic envelopes at 4 ~ 7 Hz was statistically greater during participants listening to 16-band (intelligible) than single-band (unintelligible) noise-vocoded sentences. In the study by Doelling et al. (2014), acoustic envelopes at 2 ~ 9 Hz of noise-vocoded sentences were artificially removed in various spectral bands. As a result, MEG-envelope entrainment at the corresponding rates was found to be decreased accompanied by reductions in speech intelligibility. Vanthornhout et al. (2018) used a neural reconstruction method that decodes the acoustic envelopes from EEG responses (Crosse et al., 2016) during participants recognizing speech in noisy environments. They found that the reconstruction accuracy of envelopes at 0.5 ~ 8 Hz, which reflects the degree of neural-envelope entrainment, was significantly correlated with the speech recognition performance. Moreover, association between neural entrainment and speech perception may be causally controlled by a higher-order cognitive neural network. For example, Park et al. (2015) used causal connectivity analysis in MEG showing that neural entrainment of envelopes at 1 ~ 7 Hz in intelligible (unprocessed), but not unintelligible (backward), speech, was associated with a top-down process occurring between left frontal and auditory cortices.

There have also been studies using brain stimulation, such as transcranial alternative current stimulation (tACS) that manipulated the degree of neural entrainment in order to study the causal relationship between the entrainment and speech intelligibility (Zoefel et al., 2018; Riecke et al., 2018; Wilsch et al., 2018). Zoefel et al. (2018) used tACS to manipulate phase lags between neural oscillations and the acoustic rhythm at the sentence syllable rate (~ 3 Hz). They showed that the manipulation on intelligible vocoded sentences can modulate haemodynamic responses in the superior temporal gyrus, while such findings were absent for unintelligible vocoded sentences. Riecke et al. (2018) and Wilsch et al. (2018) used similar paradigms to manipulate the neural-envelope phase lags as in Zoefel et al. (2018) (at syllable rate of 4 Hz in Riecke et al. (2018) and at the envelope rate < 10 Hz in Wilsch et al. (2018)) and found that tACS can causally modulate speech intelligibility in noisy environments.

Results of the above-mentioned studies (Peelle et al., 2013; Doelling et al., 2014; Park et al. 2015; Zoefel et al., 2018; Riecke et al., 2018; Wilsch et al., 2018; Vanthornhout et al., 2018) showed the importance of neural entrainment at the low frequencies, including delta (< 4 Hz) and theta (4 ~ 8 Hz) bands. It has been argued that entrainment at these two bands may involve different functional mechanisms (Ding and Simon, 2014). Theta-band entrainment is argued to reflect processing syllabic- and sub-syllabic-level features (Giraud and Poeppel, 2012) and it was found to covary with speech intelligibility (increased theta-band entrainment corresponding to better speech intelligibility) (Peelle et al., 2013; Ding et al., 2014). Delta-band entrainment, on the other hand, is argued to reflect processing supra-syllabic patterns such as prosodic information (Bourguignon et al., 2013; Ghitza, 2017). In contrast to theta-band entrainment, increased delta-band entrainment was found in some attention-demanding speech recognition conditions (i.e., with decreased speech intelligibility), such as recognition of speech with reduced spectral resolution (Ding et al., 2014) or with increasingly noisy backgrounds (Vander Ghinst et al., 2016). Using MEG, Molinaro and Lizarazu (2018) recently showed that delta-band, but not theta-band, entrainment is greater during processing speech than nonspeech signals in the right superior temporal and left inferior frontal regions, arguing that delta-band entrainment involves higher-order computations while theta-band entrainment is responsible for lower-level, perceptual auditory perception.

In spite of the abundant findings on the roles of neural entrainment of speech envelopes as well as distinctions between delta- and theta-band entrainment, there are still gaps with respect to linguistic and methodological concerns within these findings. First, speech intelligibility includes understanding of linguistic information at different hierarchical levels (e.g., phonology and semantics; Nahum et al., 2008). Simply seen from the relationship between neural entrainment and speech intelligibility, some critical questions still remain unanswered, e.g.: (i) What linguistic hierarchical levels are involved during the interaction between neural entrainment and speech perception? (ii) What is the role of neural entrainment and how would it subserve speech intelligibility at different hierarchical levels respectively? Second, most MEG/EEG studies reviewed above (Peelle et al., 2013; Doelling et al., 2014; Park et al. 2015; Vander Ghinst et al., 2016; Molinaro and Lizarazu, 2018; Vanthornhout et al., 2018) reported the effects of neural entrainment to single broadband acoustic envelopes. While intelligibility is achieved via human extracting acoustic components (including low-frequency envelopes) from multiple spectral bands at the cochlear output, speech with only broadband envelopes is barely intelligible (e.g., Shannon et al., 1995; Xu et al., 2005). Although speech envelopes in different spectral bands can be highly correlated with each other, such correlations reduce significantly with increased spectral distance between bands (Crouzet and Ainsworth, 2001). By applying a linear transformation algorithm on EEGs in response to speech, Di Liberto et al. (2015) provided evidence that neural encoding of envelopes from multiple spectral bands is greater than encoding of broadband envelopes. Therefore, it is important to consider that encoding multi-narrowband, rather than broadband, envelopes, could be a more appropriate form of neural entrainment. Third, although phase coherence between neural responses and acoustic envelopes (Peelle et al., 2013; Doelling et al., 2014; Vander Ghinst et al., 2016; Vanthornhout et al., 2018) provide insights into how speech acoustic features are processed, it does not characterize response functions of the brain and thus is an indirect measure of neural entrainment (Crosse et al., 2016).

By addressing these concerns from previous studies, the present study aims at characterizing the distinctions between delta- and theta-band neural entrainment at different linguistic hierarchical levels during speech perception. The present study is based on experiments and data of our previous paper which investigated the EEG oscillatory indices for different levels of auditory sentence processing (Mai et al., 2016). We used three types of continuous Mandarin utterances in order to dissociate the levels of phonology and semantics: (1) sentences consisting of meaningful disyllabic words assembled with a valid syntactic structure (‘real-word’); (2) utterances with morphologically valid syllables, but no valid disyllabic words (‘pseudo-word’); and (3) backward (time-reversed) versions of the real-word and pseudo-word utterances (for detailed descriptions, see *Stimuli and tasks* and Mai et al., 2016). Participants completed a sound-matching task when they heard an utterance in each trial and scalp-EEGs were recorded simultaneously. The types of stimuli resembled those used in previous functional imaging studies that tested the neural processing at different hierarchical levels in speech (Binder et al., 2000; Londei et al., 2010; Saur et al., 2010). Real-word and pseudo-word utterances can be distinguished by their differences in semantic contents, whilst pseudo-word and backward utterances can be distinguished by their differences in phonological contents^1^. The backward utterances were used as baselines because they are closely matched in terms of acoustic complexity to the original utterances whilst distorted phonological information (Binder et al., 2000; Londei et al., 2010; Saur et al., 2010; Gross et al., 2013). In Mai et al. (2016), we showed that several EEG signatures (band power, neural entrainment of speech envelopes, cross-frequency coupling and inter-electrode coherence) at a wide range of frequencies (delta, theta, beta and gamma) can separately index phonological and higher-level (semantic) processing. Particularly, we showed the different roles delta- and theta-band neural entrainment, where the theta-band entrainment indexes greater phonological processing for speech (real-word and pseudo-word) than for non-speech (backward) while delta-band entrainment indexes greater effortful phonological recognition for pseudo-word utterances. However, similar to previous studies (e.g., Peelle et al., 2013; Doelling et al., 2014; Vander Ghinst et al., 2016), phase coherence between EEGs and the speech broadband envelopes, an indirect measure of neural entrainment, was calculated. In the present study, neural entrainment was quantified using a linear transformation algorithm via multivariate Temporal Response Functions (mTRF) (Di Liberto et al. 2015; Crosse et al., 2016). Such approach characterizes the brain’s response function that maps acoustic features onto neural responses, providing a more direct measure of neural entrainment. It can also reflect EEG encoding of multi-narrowband envelopes (see details in Crosse et al., 2016, and *Methods*), outweighing measures of neural entrainment to broadband envelopes in many other studies. With the syllable rate of all utterances being controlled at around 4 Hz, delta- and theta-band were defined as 1.5 ~ 3 Hz (average cycle at 500 ms corresponding to 2 Hz) and 3 ~ 6 Hz (average cycle at 250 ms corresponding to 4 Hz), respectively. Delta- and theta-band thus respectively corresponded to rhythms at supra-syllable and syllable/sub-syllable rates.

We hypothesize that, due to delta- and theta-band neural entrainment reflecting processing of speech at different cognitive stages (Ding and Simon, 2014), they should also take distinct roles at different linguistic hierarchical levels. Particularly, as theta rhythms were argued to reflect the tracking of syllabic and sub-syllabic information (Peña and Melloni, 2012; Giraud and Poeppel, 2012) that convey phonological contents (Rimol et al., 2005), we predict that theta-band entrainment should be involved in phonological processing. On the other hand, as delta-band entrainment may be related to higher-order cognitive processing (Ding et al., 2014; Vander Ghinst et al., 2016; Molinaro and Lizarazu, 2018), we predict that delta-band entrainment is involved in semantic-level processing. To test such hypotheses, delta- and theta-band entrainment were measured and compared statistically across the stimulus types (real-word, pseudo-word and backward utterances). Subsequently, capacities of mTRFs on classifying EEG trials into correct stimulus types were further tested to determine the specificity of neural entrainment at different hierarchical levels. Temporal properties of mTRFs were finally examined to study how the degrees of delta- and theta-band entrainment vary across the timescales of neural processing for different stimulus types. We suggest that testing our hypotheses will consolidate our understanding on neural entrainment of low-frequency envelopes during speech perception.

## 2. Methods

The present study used the EEG data collected from our previous study that investigated the relationship between brain oscillations and auditory sentence processing (Mai et al., 2016). Participants, stimuli and experiment paradigms had all been previously described in this study.

### 2.1 Participants

Twenty normal-hearing, native Mandarin speakers from mainland China (8 male; aged 19 ~ 25 years old) were recruited and paid for participating the experiment. No history of neurological disorders were reported for any participant. All participants were either right-handed (18 participants with handedness indices (HI) > 40) or towards right-handed (2 participants with HIs = 33.3) according to the Edinburgh Handedness Inventory (Oldfield, 1971).

### 2.2 Stimuli and tasks

Stimuli consisted of three types of continuous Mandarin utterances: (1) real-word, (2) pseudo-word, and (3) backward utterances. (1) and (2) were naturally produced by a male native Mandarin speaker recorded at a sampling rate of 22,050 Hz. All were produced with syllable rates between 3.5 and 4.5 Hz, and some were adjusted by slightly lengthening or shortening in time via software PRAAT (University of Amsterdam, The Netherlands) in order to keep all utterances at ~ 4 Hz syllable rate. The real-word utterances were semantically unpredictable sentences (SUSs) (Benoit et al., 1996). Each SUS here was comprised of four semantically valid disyllabic (two-character) words with a syntactic structure of ‘Subject + Verb + Attribute + 的 + Object’. Character ‘的’ is a grammatical particle without lexical meaning. The words within a sentence were not contextually related to each other and it was impossible to predict a word from the sentence it is in. A sample SUS was ‘网络喜欢坚强的 空气’, in which the disyllabic words were ‘网络’ (‘Internet’), ‘喜欢’ (‘enjoy’), ‘坚强’ (‘tough’), and ‘空气’ (‘air’). The purpose of using SUSs was to prevent participants from identifying sentence contents from contextual information and to guarantee that they attended to the entire utterance. Pseudo-word utterances were sentences consisting of the same number of morphologically valid syllables as in each real-word utterance, but with no two adjacent syllables forming a semantically valid word. All participants confirmed that all pseudo-word utterances were semantically invalid for them after the experiment. Backward utterances were time-reversed versions of the real-word and pseudo-word utterances, which caused substantial phonological distortion but retain similar acoustic complexity of the speech (temporal fluctuations, formant distributions, and harmonic structures) (Binder et al., 2000; Londei et al., 2010; Saur et al., 2010; Gross et al., 2013).

There were 80 utterances for each of the three stimulus types without repetition of any utterance. Half of the backward utterances were generated from randomly selected real-word utterances with the other half from randomly selected pseudo-word utterances. All stimuli had a similar duration (2.2 ~ 2.3 seconds) and were adjusted to the same average RMS intensity.

During the experiments, participants were seated in front of a computer screen and listened to the stimuli via EARTONE 3A inserted earphones (Etymotic Research, USA) with a fixed loudness at ~ 70 dB for all utterances. All stimuli (three types with 80 trials for each type) were presented in a random order using EPrime 2.0 (Psychology Software Tools, USA). The paradigm of each trial is shown in **Fig. 1**. At the start of each trial, there was a 3-second silence allowing participants to blink, followed by another 1.5-second silence with a white cross centred on the screen. A cue sound (200 ~ 300 ms; a naturally produced syllable for the real-word and pseudo-word utterances, or a backward syllable for the backward utterances) was then presented. These were then followed by a 2-second silence and the target utterance. Participants were required to complete a sound-matching task. They were instructed to make a forced-choice judgement whether the cue sound was present in the target utterance or not by pressing a button representing ‘Yes’ or ‘No’ (on the left or right side of the keyboard) when a question mark appeared on the screen after the utterance. They were instructed to sit still, keep their eyes on the white cross and avoid any eye blink or body movement after the cue sound was played. They were also asked to press the button *only* after the question mark appeared, in order to avoid motor artefacts during the target period. Feedback of accuracies was given every 30 trials and participants were encouraged to respond as accurately as possible. Overall, the aim of the sound-matching task was to keep participants actively attending to the target utterances.

**Fig. 1.**
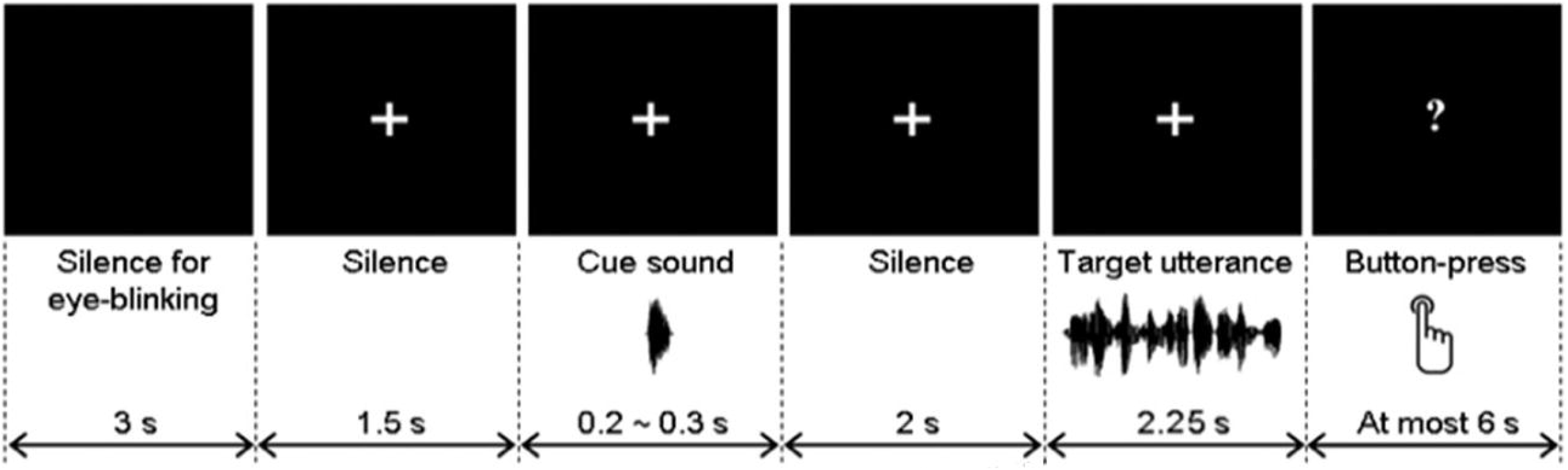
Time course of each trial in the experiment. Visual presentations were shown as the top panels and texts at the bottom describe the corresponding time course of the audio presentation and the sound-matching task (button-press). The figure is adopted from Mai et al. (2016) with permission.

Out of all 80 trials in each stimulus type, 20% of the trials in which the cue sounds were actually present in the target utterances (16 utterances). In the present study, only the trials where the cue sounds were *not* present in the target utterances (64 utterances) were included in the subsequent analyses. This was to preclude the possibility of participants not attending to the entire utterance period and to avoid auditory repetition effects when the cue sound was present in the target utterance. This could also minimize possible effects of motor preparation of button press due to judgments made before the end of the utterance when the cue sound was present.

30 practice trials (utterances all different from the formal test) were run prior to the formal test. Breaks were taken every 30 trials during the formal test.

### 2.3 EEG recording and preprocessing

Scalp-EEGs were recorded via a 32-electrode ActiveTwo system (Biosemi, The Netherlands) sampled at 1024 Hz. Bilateral mastoids were used as the reference. Eye artefacts were detected via vertical (vEOG; electrodes above and below the left eye) and horizontal EOGs (hEOG; electrodes on the lateral sides of the left and right eyes).

Signals of all electrodes (including EOGs) were first re-referenced to the bilateral mastoids and then bandpass filtered at 0.5 ~ 8 Hz using a zero-phase, 2nd-order Butterworth filter. Signals for detecting eye artefacts were then obtained by subtracting between signals in corresponding EOG electrodes (vEOGs and hEOGs for vertical and horizontal artefacts, respectively). Trials where the filtered EEGs in the target period (target utterances with a fixed length of 2.25 seconds for all trials) exceeded ± 40 μV in any electrode (including vEOG and hEOG) were treated as being contaminated by eye or body movement artefacts and were rejected from subsequent analyses.

### 2.4 Extraction of delta- and theta-band EEGs and stimulus envelopes

Delta- and theta-band neural entrainment were calculated via a linear transformation algorithm based on multivariate Temporal Response Functions (mTRF) (Di Liberto et al., 2015; Crosse et al., 2016). The algorithm calculates the extent of mapping speech envelope information onto corresponding EEG responses. The algorithm was applied on delta- and theta-band entrainment separately for the three stimulus types (real-word, pseudo-word and backward) in each participant. Delta-band entrainment was quantified based on the delta-band EEGs and stimulus envelopes, whilst theta-band entrainment was quantified based on the theta-band EEGs and stimulus envelopes.

EEGs were bandpass filtered at 1.5 ~ 3 Hz (delta) and 3 ~ 6 Hz (theta) using a zero-phase, 2nd-order Butterworth filter. The signals were then decimated to 128 Hz via a 30th-order Hamming-windowed FIR filter. The delta- and theta-band EEG signals within the artefact-free target periods were then respectively used for quantifying delta- and theta-band entrainment.

The stimulus acoustic envelopes of the artefact-free trials were obtained as follows. First, each corresponding utterance was bandpass filtered between 100 and 5000 Hz and then resampled to 16384 Hz (an integer multiple of 128 Hz) using PRAAT. Second, delta- and theta-band envelopes of each utterance was extracted based on either a single broadband (‘BROAD’) or multiple narrowbands (‘MULTI’). For the BROAD condition, delta- and theta-band envelopes were obtained by bandpass filtering the broadband Hilbert envelope of the utterance at 1.5 ~ 3 Hz and at 3 ~ 6 Hz (using the same filter as in EEGs), respectively. For the ‘MULTI’ condition, the utterance were bandpass filtered into 16 logarithmic-spaced acoustic spectral bands between 100 and 5000 Hz. The delta- and theta-band envelopes were then extracted from each spectral band following the same way as in the ‘BROAD’ condition. All acoustic envelopes were finally decimated to 128 Hz as in EEGs. In this way, for both delta- and theta-band envelopes, there was only one envelope time series in the ‘BROAD’ condition, but 16 envelope time series in the ‘MULTI’ condition.

### 2.5 Calculations of TRFs

Temporal Response Functions (TRFs) (Di Liberto et al., 2015; Crosse et al., 2016) for all artefact-free trials were then estimated using a linear transformation algorithm:

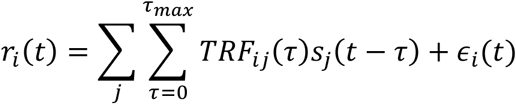

Where *i* and *j* refer to the ith electrode and the *j*th spectral band of the acoustic stimulus, respectively; *r_i_*(*t*) is the EEG time series; *TRF_ij_*(*t*) is the time series of the TRF; *s_j_*(*t*) is the time series of the stimulus envelopes; *ϵ_i_*(*t*) is the normally-distributed error term; τ_max_ is the maximum time lag between the EEG series and the stimulus series, which was set at 300 ms in the present study. The *TRF_ij_*(*t*) was estimated by minimizing the mean squares of *ϵ_i_*(*t*). As such, TRF can be obtained via the following matrix formula:

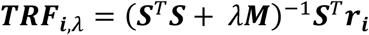

where *i* refers to the *i*th electrode; ***S*** is a matrix comprised of lagged time series of the stimulus envelopes in all spectral bands; ***r**_i_* is the vector of EEG series; *λ* and ***M*** denote the ridge regression parameter and a quadratic matrix, respectively, during the regularization that avoided the ill-posed estimation and overfitting (see Crosse et al., 2016). The ridge regression parameter *λ* was chosen among a range of values (2^−15^, 2^−14^,…, 2^14^, 2^15^) and the optimal *λ* was obtained according to the cross-validation during the training stage (see *Training and testing*).

### 2.6 Training and testing

Artefact-free trials were divided into a training set and a testing set during the procedure of training and testing. Here, for each stimulus type (real-word, pseudo-word or backward) and each participant, we randomly assigned 50 trials to the training set and randomly selected one of the remaining trials as the testing trial. We replicated the training and testing procedure for 1000 times and the final testing result was treated as the average over the corresponding 1000 testing estimates (‘predictive powers’ or *PredPowers*, see below). We followed this procedure due to the different numbers of artefact-free trials across stimulus types and across participants (recall that there were 64 trials for each stimulus type *prior to* artefact rejection). We considered that this procedure could keep the number of training and testing trials (50 and 1, respectively) the same for all stimulus types and participants, and at the same time all trials had similar chances to be either trained or tested.

During the training stage, a ‘leave-one-out’ cross-validation procedure was followed in order to obtain the optimal ridge parameter *λ* and the trained TRF (Crosse et al., 2016). First, in the training set, one trial was chosen to be ‘left out’ as a validator, while the remaining trials were treated as a ‘sub-training’ set. A predictive EEG series was generated for the validator using the temporal average of the TRFs across the trials in the sub-training set:

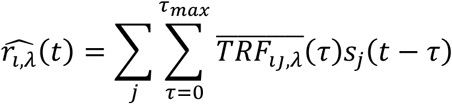

where 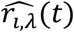 is the predictive EEG series; 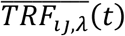 is the average TRF series across the trials in the sub-training set; and *s_j_*(*t*) is the stimulus envelope of the validator. The Pearson correlation (Fisher-transformed) and the mean-squared error (MSE) between 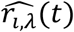 and the actual EEG series of the validator were calculated. Second, a different trial was then selected as the validator in the next round of validation. The validation procedure was repeated until all trials in the training set were assigned as validators. The correlation values and MSEs were then averaged across all validators. The optimal *λ* value was identified as the one which yielded the highest correlation value or the lowest MSE. In the present study, we used *λ* which yielded the highest correlation value but not the lowest MSE, as we found that the predictive powers were significantly greater based on the former than on the later (see *Results*). The trained TRF was finally obtained by averaging the TRFs with the optimal *λ* value across all trials in the training set.

During the testing stage, the predictive EEG series was obtained based on the trained TRF and the stimulus envelope series of the testing trial, following the same procedure as in each round of validation in the training stage. Then the Pearson correlation was calculated between the predictive EEG series and the actual EEG series of the testing trial. The ‘predictive power’ (*PredPower*) was quantified as the Fisher-transform of the correlation value. The final *PredPower* was obtained as the average across the 1000 times of training and testing.

### 2.7 Surrogate and random predictive powers (*PredPowers*)

Surrogate and random *PredPowers* were calculated as baselines to prove the fidelity of the results obtained from our data. Surrogate *PredPowers* were obtained as follows. During each round of training and testing, the testing trial selected for each given stimulus type was assigned to the other two stimulus types as a testing ‘surrogate’. The surrogate *PredPowers* for a given stimulus type were then defined as the predictive powers that were obtained using testing trials from different stimulus types. The results were finally averaged across all stimulus types and across the 1000 times of training and testing. To obtain random *PredPowers*, 51 random ‘trials’ (50 for the training set and one for testing) were created, each of which consisted of an ‘EEG’ series and a corresponding ‘stimulus’ series, both being pseudo-randomly generated Gaussian noises with the same length of each target period in the experiment (2.25 seconds). *PredPowers* were calculated in the same way as in the real data. Such procedure was replicated 1000 times and the random *PredPowers* were finally grand-averaged.

We predicted that, if *PredPowers* are valid measurements and TRFs can specifically encode envelope information of the respective stimulus types, the ‘congruent’ *PredPowers* (those based on testing trials from the same stimulus type) should be statistically greater than both surrogate and random *PredPowers*. We also predicted that, as some acoustic features (such as acoustic rhythms and spectrotemporal complexity) are commonly shared between stimulus types, surrogate *PredPowers* could also be above random level.

### 2.8 Classification of EEG trials using mTRFs

Classification capacity was tested for multivariate TRFs (mTRFs). If *PredPowers* can index the neural entrainment at different linguistic hierarchical levels, mTRFs should have the capacity to classify EEGs between different stimulus types. Similar to the calculation of surrogate *PredPowers*, the testing trial in each stimulus type was assigned to the other two stimulus types as a testing surrogate during each round of training and testing. As such, the mTRF in each given stimulus type generated one congruent *PredPower* and two surrogate *PredPowers*. The capacity of mTRF was estimated by whether it could accurately identify the congruent testing trial (the trial from the same stimulus type). We considered that the classification of mTRF was ‘accurate’ if the congruent *PredPower* was greater than the surrogate *PredPowers*. The accuracies were finally averaged over the 1000 times of training and testing.

Furthermore, the classification capacity in each stimulus type was estimated in two scenarios: (1) classification among all stimulus types, i.e., when mTRF of each given stimulus type was tested by trials from all three stimulus types (Scenario_1); (2) classification between two stimulus types, i.e., when mTRF of each given stimulus type trials was tested by trials from two stimulus types, one from the same stimulus type of the given mTRF and the other from a different stimulus type (e.g., mTRF_real-word_ with testing trials from real-word and pseudo-word utterances) (Scenario_2).

### 2.9 Temporal properties of mTRFs

Time series of mTRF were obtained by averaging over the 1000 trained mTRFs for each stimulus type and each participant. Absolute values of the time series were then obtained as the absolute weighting series in each spectral band. Absolute values were used here because they could reflect the extent of mTRF contributions to the neural entrainment regardless of the sign of the weighting. The absolute series were averaged across all spectral bands (i.e., 16 bands).

Temporal properties of the absolute weighting were then examined to study how the degrees of neural entrainment vary across time lags. Recall that the range of time lags was set as 0 ~ 300 ms (see *Calculations of TRFs*). The time lags were divided into ‘early’ (20 ~ 160 ms) and ‘later’ (160 ~ 300 ms) stages (each covered 140 ms). The absolute weighting were compared across the time ranges (‘early’ vs. ‘later’) and stimulus types.

### 2.10 Sequences of statistical analyses

Calculations of *PredPowers* and TRFs were all electrode-wise and separately conducted for delta- and theta-band entrainment, based on the delta- and theta-band EEGs and stimulus envelopes, respectively (see *Extraction of delta- and theta-band EEGs and stimulus envelopes*). Statistical analyses were also conducted for delta- and theta-band entrainment separately. Also, the analyses were conducted based on *PredPowers* and TRFs averaged over the centro-frontal electrodes. This is because the neural entrainment measured with EEGs is dominant over centro-frontal (compared to parieto-occipital) region for the auditory modality (Crosse et al., 2015, 2016). The centro-frontal electrodes were defined as the 22 electrodes shown in **Fig. 2** (indicated by the shaded trapezoid). All statistical analyses were within-subject analyses (Repeated Measures ANOVAs followed by post-hoc pairwise t-tests).

**Fig. 2.**
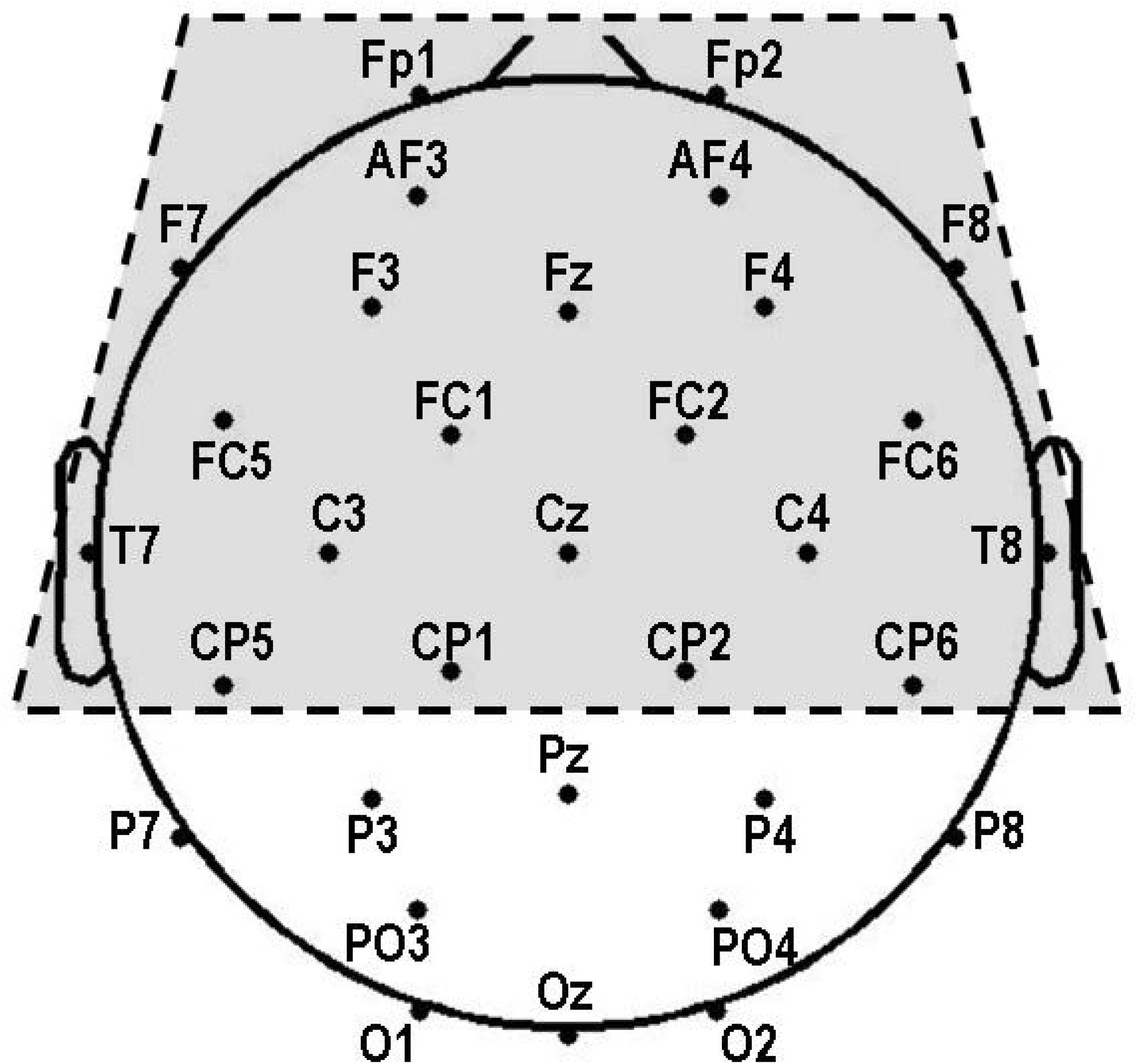
Channel configuration. Statistical analyses were based on the centro-frontal electrodes (indicated by the shaded trapezoid).

*PredPowers* were first compared between the ‘BROAD’ condition (using TRFs based on the broadband envelopes of the stimuli, or univariate TRFs) and the ‘MULTI’ condition (using TRFs based on stimulus envelopes extracted from 16 spectral bands, or multivariate TRFs (mTRFs)) (see *Extraction of delta- and theta-band EEGs and stimulus envelopes*). Results showed that *PredPowers* were significantly greater in the ‘MULTI’ than in the ‘BROAD’ condition at both delta and theta bands (see *Results*). This is consistent with previous findings showing that mTRF models are superior to univariate TRF models for predicting low-frequency EEG responses during speech perception (Di Liberto et al., 2015). Accordingly, we used the mTRF, but not univariate TRF model, during the subsequent signal processing and statistical analyses.

Fidelity of *PredPowers* were then tested by comparing those with the surrogate and random *PredPowers*. Next, *PredPowers* were compared across the three stimulus types (real-word, pseudo-word and backward) to test how neural entrainment changes at different linguistic hierarchical levels. Classification capacity of mTFRs were then tested in order to examine the specificity of neural entrainment for different stimulus types. Temporal properties of mTRFs were finally examined to study the degrees of neural entrainment across time lags.

The EEG signal processing was conducted using Matlab 2014a (MathWorks, USA). Statistical analyses were conducted using SPSS 23 (IBM, USA).

## 3. Results

At least 51 trials were retained after artefact rejection in all stimulus types for all participants. The average numbers of retained trials were 59.1 (SE: 0.7), 58.3 (SE: 0.7) and 58.9 (SE: 0.9) for real-word, pseudo-word and backward utterances, respectively. No significant difference for the number of trials was found between any two stimulus types (all *p* > 0.1, uncorrected). Behavioral results can be found in Mai et al (2016). Response accuracies were significantly higher than the 50% chance-level for all stimulus types (> 95% for real-word and pseudo-word utterances and > 70 % for the backward utterances; all *p* < 10^−8^, uncorrected), indicating that participants had complied with the instructions to actively attend to the stimuli.

All statistical analyses on EEGs were conducted based on the averages over the 22 centro-frontal electrodes (see *Methods*). Repeated Measures ANOVAs were conducted with Greenhouse-Geisser correction. All *p*-values in the pairwise comparisons between any two stimulus types were Bonferroni corrected by the factor of 3 (due to the three stimulus types) unless specified as ‘uncorrected’.

### 3.1 Univariate TRF vs. mTRF

*PredPowers* were compared between the univariate TRF and mTRF models. Before such comparisons were conducted, it was first determined that the optimal *λ* value (the ridge regression parameter, see *Methods*) was identified as the one which yielded the highest Pearson correlation value (Fisher-transformed) but not the lowest MSE during the cross-validation. This was because *PredPowers* were found to be significantly greater based on the former than on the latter in both univariate TRF and mTRF models (all *p* < 0.01).

Repeated Measures ANOVAs were then conducted for the delta- and theta-band *PredPowers* with factors of TRF Type (univariate TRF vs. mTRF) and Stimulus Type (real-word vs. pseudo-word vs. backward). The results showed significant main effects of TRF Type and Stimulus Type, but no [TRF Type × Stimulus Type] interactions, for both delta- and theta-band *PredPowers* (see **Table 1** for detailed statistics). Specifically, both delta- and theta-band *PredPowers* were significantly greater when using mTRF compared to univariate TRF (see **Fig. 3**). As we only focused on the differences of the two TRF types in this section, post-hoc analyses following the main effects of Stimulus Type are not reported here.

**Fig. 3.**
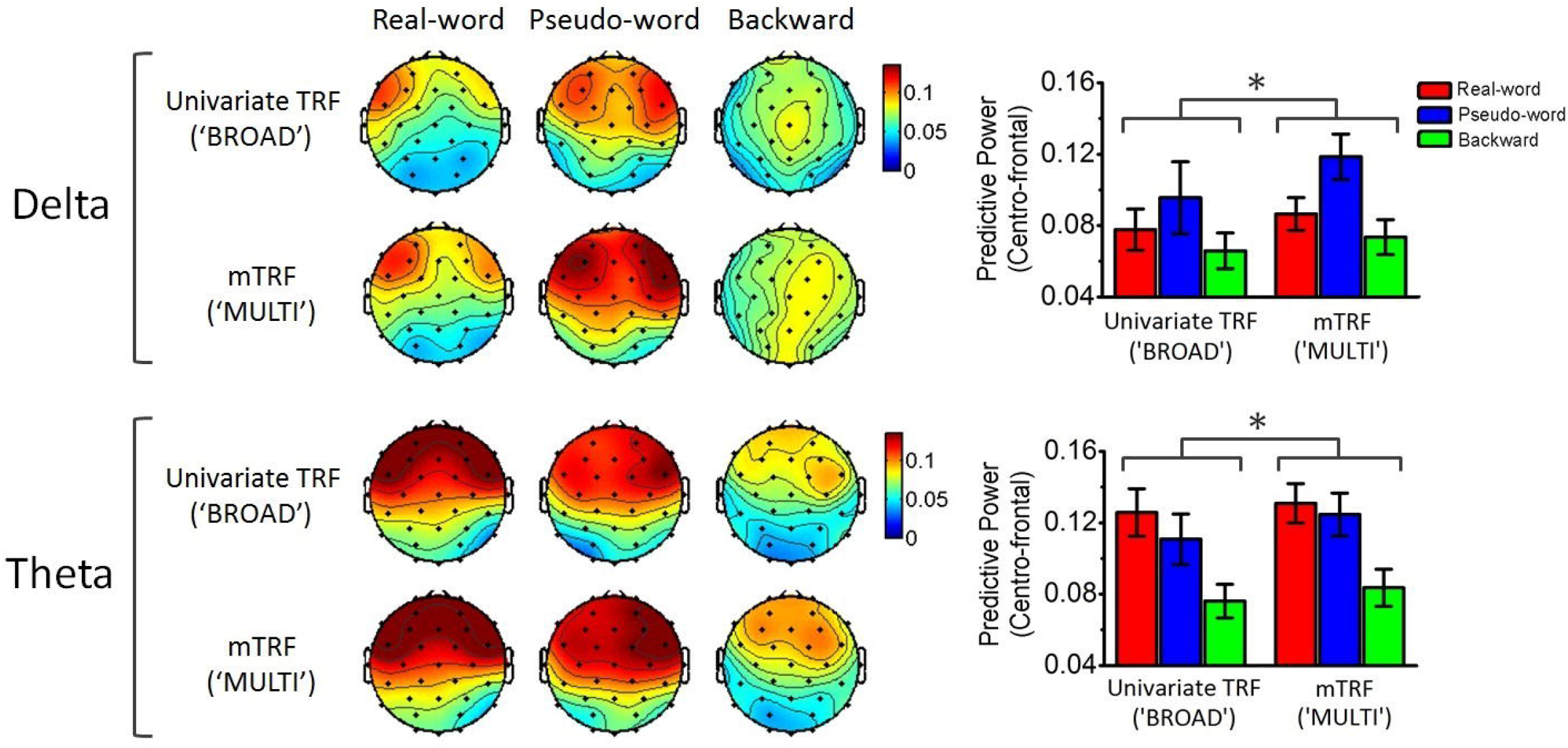
Comparisons of *PredPowers* between univariate TRF (‘BROAD’) and mTRF (‘MULTI’) models. Scalp topographies for different stimulus types are shown on the left and *PredPowers* averaged across the centro-frontal electrodes were shown on the right. Errors bars denote standard errors of the mean (SEMs). * = significance at *p* < 0.05.

**Table 1.** Statistical results of Repeated Measures ANOVAs for *PredPowers* averaged over the centro-frontal electrodes. The effects of the TRF type (univariate TRF vs. mTRF) and Congruency (congruent vs. surrogate; for *PredPowers* based on mTRF) were tested, respectively. *Df, F, p* and *η_p_^2^* refer to degrees of freedom, F-values, *p*-values and partial eta-squared, respectively. The statistics were Greenhouse-Geisser corrected. Numbers are all rounded to three decimal places, unless they are < 0.001. Significant *p*-values are indicated in bold. * = significance at *p* < 0.05; ** = significance at *p* < 0.01; *** = significance at *p* < 0.001.

| Dependent variables | Band | Factors | *df1* | *df2* | *F* | *p* | $\eta_p^2$ |
| --- | --- | --- | --- | --- | --- | --- | --- |
| *PredPower* | Delta | TRF Type | 1 | 19 | 5.369 | **0.032\*** | 0.220 |
|  |  | Stimulus Type | 1.710 | 32.494 | 5.429 | **0.012\*** | 0.222 |
| | | TRF Type × Stimulus Type | 1.369 | 26.003 | 1.196 | 0.302 | 0.059 |
|  | Theta | TRF Type | 1 | 19 | 6.650 | **0.018\*** | 0.259 |
|  |  | Stimulus Type | 1.539 | 29.233 | 7.543 | **0.004\*\*** | 0.284 |
| | | TRF Type × Stimulus Type | 1.759 | 33.424 | 1.290 | 0.286 | 0.064 |
| *PredPower* based on mTRF | Delta | Congruency | 1 | 19 | 26.638 | **$< 10^{-4}$\*\*\*** | 0.584 |
|  |  | Stimulus Type | 1.950 | 37.056 | 7.510 | **0.002\*\*** | 0.283 |
| | | Congruency × Stimulus Type | 1.830 | 34.769 | 1.196 | **0.003\*\*** | 0.282 |
|  | Theta | Congruency | 1 | 19 | 40.280 | **$< 10^{-5}$\*\*\*** | 0.679 |
|  |  | Stimulus Type | 1.313 | 24.944 | 7.602 | **0.007\*\*** | 0.286 |
| | | Congruency × Stimulus Type | 1.710 | 32.486 | 4.746 | **0.020\*** | 0.200 |

The results thus showed the superiority of mTRF to univariate TRF, consistent with the previous finding (Di Liberto et al., 2015). The subsequent signal processing and statistical analyses were hence based on mTRF, but not on univariate TRF.

### 3.2 Fidelity of *PredPowers*

Fidelity of *PredPowers* (based on mTRFs) were tested by comparing congruent *PredPowers* (training and testing trials from the same stimulus type) with surrogate (training and testing trials from different stimulus types) and random (pseudo-random noise) *PredPowers*. The results are illustrated in **Fig. 4**.

**Fig. 4.**
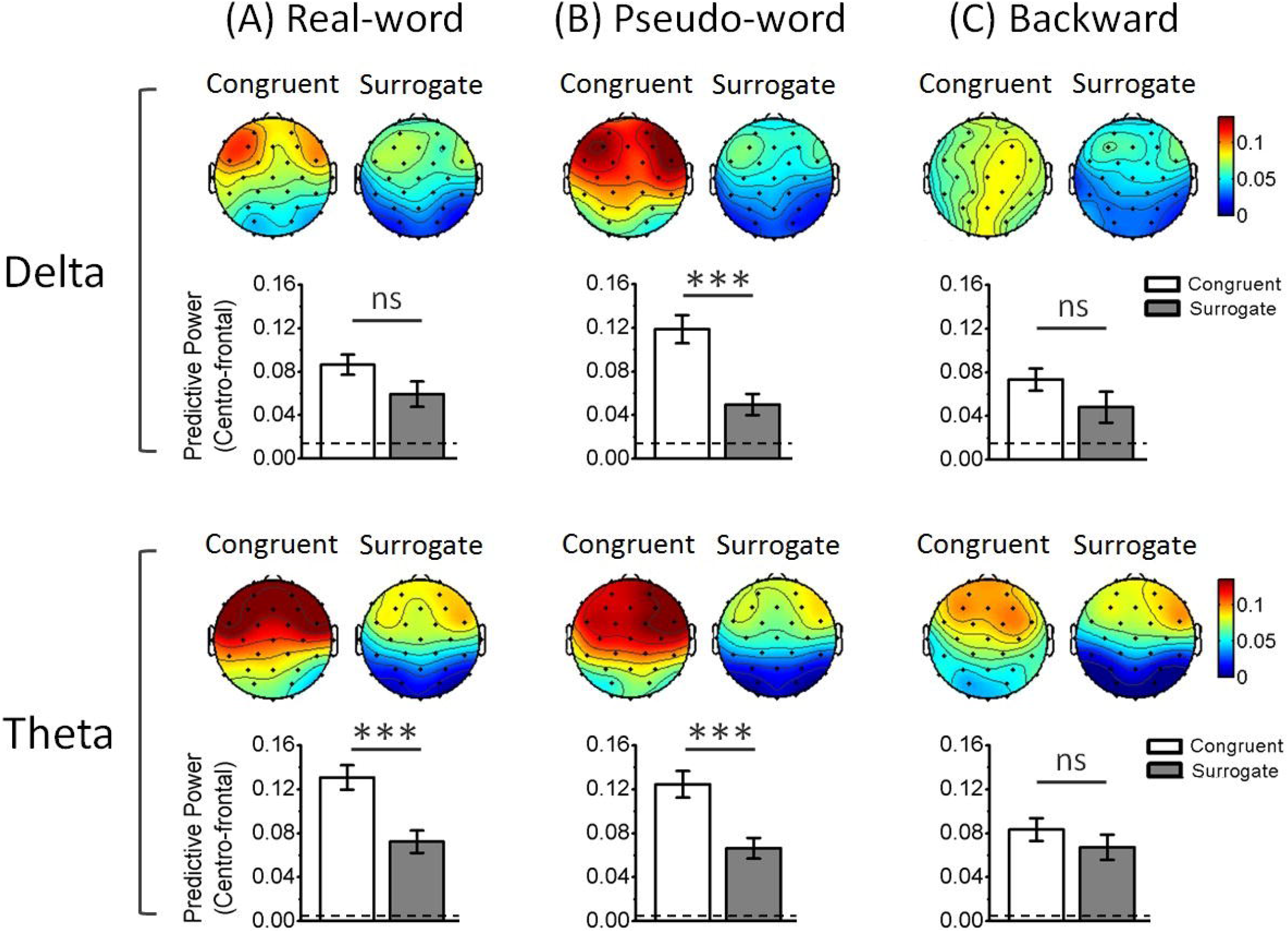
Comparisons between congruent and surrogate *PredPowers*. All *PredPowers* were calculated based on the mTRF model. Bar graphs illustrated the comparisons averaged over the centro-frontal electrodes for different stimulus types. Statistical significance were Bonferroni corrected by the factor of 3 (three stimulus types). Dashed lines indicate the values of random *PredPowers*. Errors bars denote SEMs. *** = significance at *p* < 0.001; ns = not significant.

We first test whether *PredPowers* obtained from real data (congruent and surrogate) were statistically above random level. We found that all congruent *PredPowers* at delta and theta bands were significantly greater than random *PredPowers* for all stimulus types (all *p* < 10^−4^, uncorrected). Surrogate *PredPowers* were significantly greater than random *PredPowers* for all stimulus types (all *p* < 0.005, uncorrected), except that at the delta-band for the backward utterances (*p* = 0.052, uncorrected). We suggest it is reasonable that, not only congruent *PredPowers*, but also surrogate *PredPowers* were greater than the random level, possibly because some acoustic features (e.g., acoustic rhythms and spectrotemporal complexity) were commonly shared across different stimulus types, resulting in these features being encoded in mTRFs.

Next, the effects of Congruency (congruent vs. surrogate) were tested. Repeated Measures ANOVAs were conducted with factors of Congruency and Stimulus Type. Main effects of Congruency and Stimulus Type, and [Congruency × Stimulus Type] interactions were all found to be significant at both delta and theta bands (see **Table 1**). Post-hoc pairwise comparisons following the significant interactions showed that, at the delta band, congruent *PredPower* was significantly greater than surrogate *PredPower* only for pseudo-word utterances (t_(19)_ = 7.984, *p* < 10^−6^), but not for real-word (t_(19)_ = 2.449, *p* = 0.073) or backward utterances (t_(19)_ = 2.056, *p* = 0.161) (see **Fig. 4** upper panels). At the theta band, congruent *PredPower* was significantly greater than surrogate *PredPower* for real-word (t_(19)_ = 5.769, *p* < 10^−5^) and pseudo-word utterances (t_(19)_ = 5.733, *p* < 10^−5^), but not for backward utterances (t_(19)_ = 1.162, *p* = 0.779) (see **Fig. 4** lower panels).

### 3.3 Comparisons of *PredPowers* between stimulus types

Results for comparisons of *PredPowers* between stimulus types are illustrated in **Fig. 5**. Repeated Measures ANOVAs were conducted with the factor of Stimulus Type. Significant main effects of Stimulus Type were found for both delta- and theta-band *PredPowers* (see **Table 2**). Post-hoc comparisons showed that, at the delta band, *PredPower* was significantly greater for pseudo-word than for real-word and backward utterances, while no significant difference was found between real-word and backward utterances (see **Fig. 5** and **Table 3**, delta-band *PredPowers*). At the theta band, *PredPower* was significantly greater for real-word and pseudo-word than for backward utterances, while no significant difference was found between real-word and pseudo-word utterances (see **Fig. 5** and **Table 3**, theta-band *PredPowers*).

**Fig. 5.**
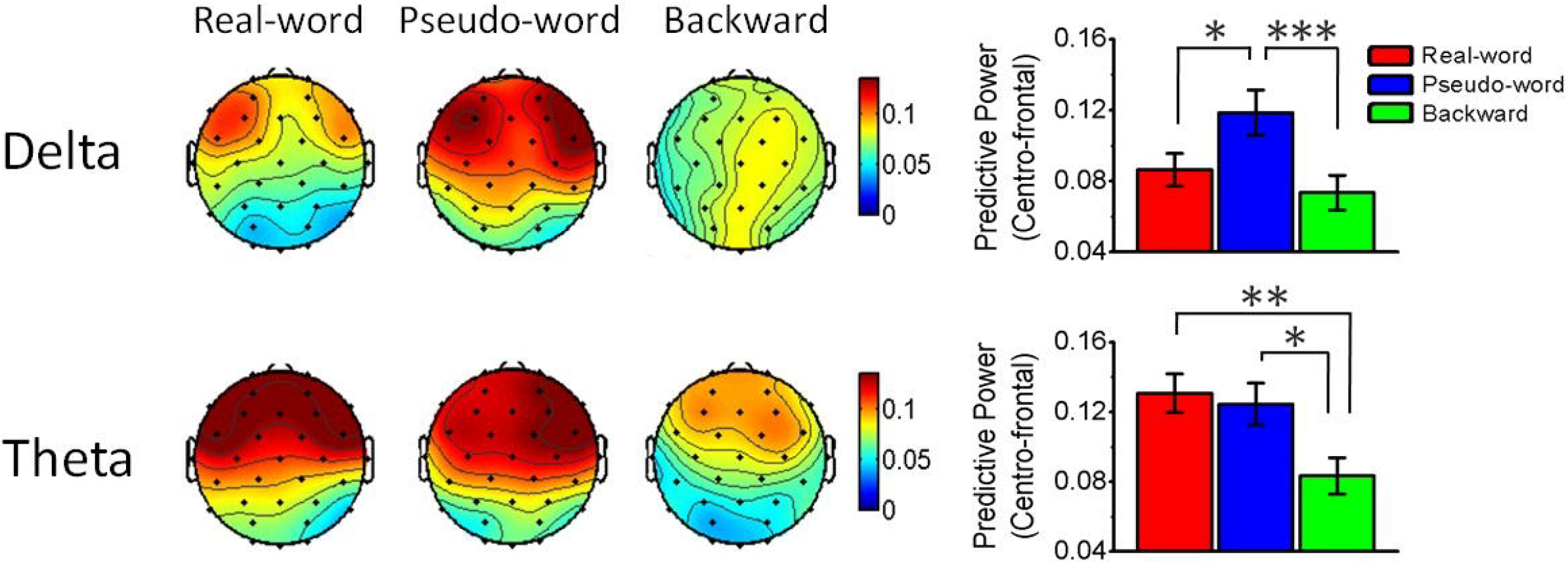
Comparisons of *PredPowers* across stimulus types. All *PredPowers* were calculated based on the mTRF model. Bar graphs illustrated the comparisons averaged over the centro-frontal electrodes. Statistical significance were Bonferroni corrected by the factor of 3. Errors bars denote SEMs. * = significance at *p* < 0.05; ** = significance at *p* < 0.01; *** = significance at *p* < 0.001.

**Table 2.** Statistical results of Repeated Measures ANOVAs for *PredPowers* (based on mTRF) and the classification capacity of mTRFs across stimulus types. All were based on the centro-frontal electrodes. Note that ANOVAs for the mTRF classification were conducted only in Scenario_1 (when mTRFs were tested by trials from all three stimulus types), but not in Scenario_2 (when mTRFs were tested by trials from two stimulus types). *Df, F, p* and *η_p_^2^* refer to degrees of freedom, F-values, *p*-values and partial eta-squared, respectively. The statistics were Greenhouse-Geisser corrected. Numbers are all rounded to three decimal places, unless they are < 0.001. Significant p-values are indicated in bold. * = significance at *p* < 0.05; ** = significance at *p* < 0.01; *** = significance at *p* < 0.001.

| Dependent variables | Band | Factors | *df1* | *df2* | *F* | *p* | $\eta_p^2$ |
| --- | --- | --- | --- | --- | --- | --- | --- |
| *PredPower* based on mTRF | Delta | Stimulus Type | 1.633 | 31.019 | 11.105 | **< 0.001\*\*\*** | 0.369 |
|  | Theta | Stimulus Type | 1.555 | 29.542 | 7.956 | **0.003\*\*** | 0.295 |
| Classification accuracy of mTRF (Scenario_1) | Delta | Stimulus Type | 1.823 | 31.019 | 6.793 | **0.004\*\*** | 0.263 |
|  | Theta | Stimulus Type | 1.902 | 36.133 | 3.344 | **0.049\*** | 0.150 |

**Table 3.**
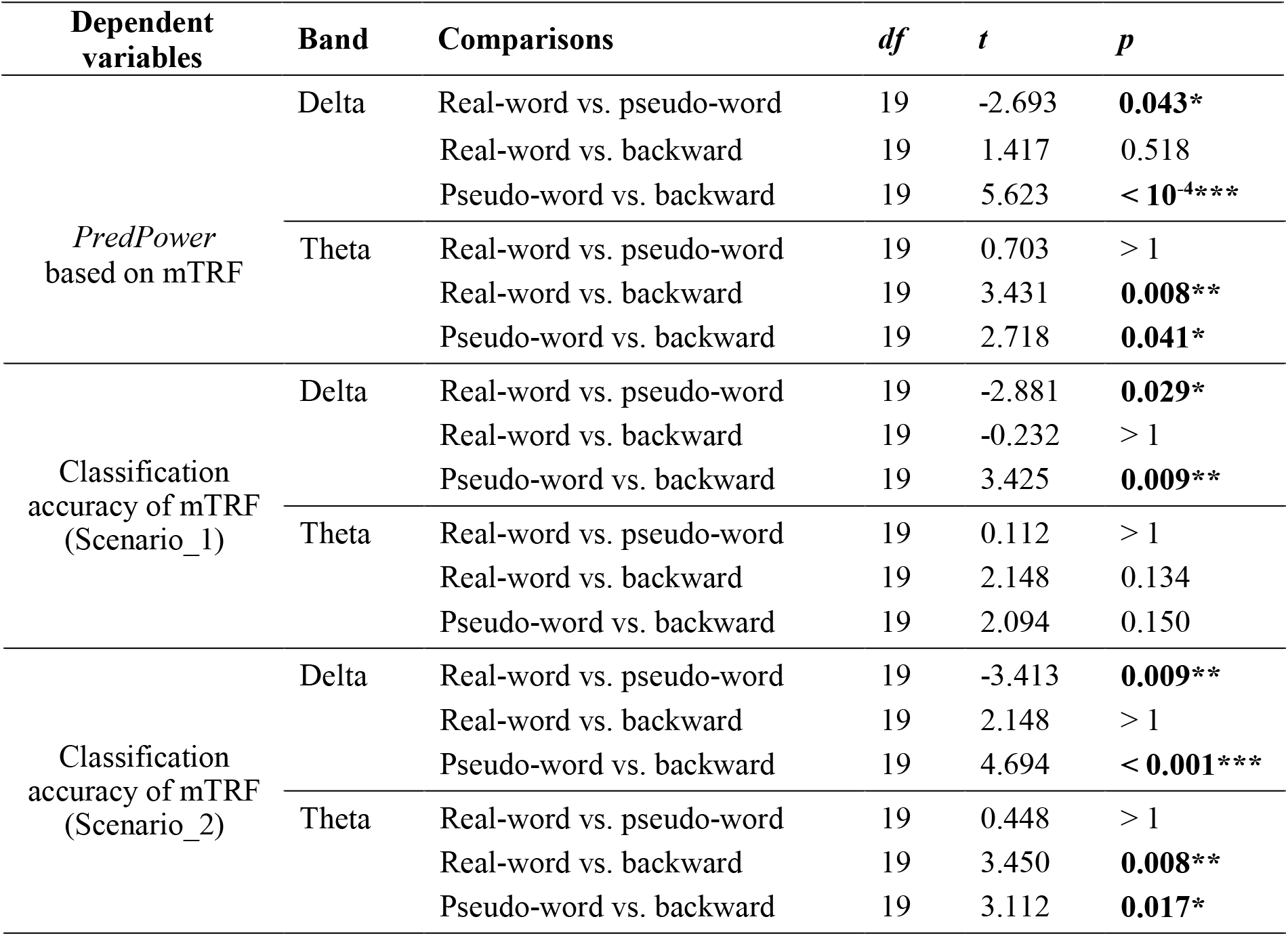
Pairwise comparisons for *PredPowers* (based on mTRF) and the classification capacity of mTRFs between different stimulus types. The comparisons for *PredPowers* and classification accuracies of mTRFs in Scenario_1 were post-hoc analyses following the significant main effects of Stimulus Type (see **Table 2**). *Df, t* and *p* refer to degrees of freedom, t-values and *p*-values, respectively. All *p*-values were Bonferroni corrected by the factor of 3 (three stimulus types). Numbers are all rounded to three decimal places, unless they are < 0.001 or *p* > 1. Significant *p*-values are indicated in bold. * = significance at *p* < 0.05; ** = significance at *p* < 0.01; *** = significance at *p* < 0.001.

### 3.4 Classification capacity of mTRFs

**Fig. 6** and **Fig. 7** show the results of classification capacity of mTRFs.

**Fig. 6.**
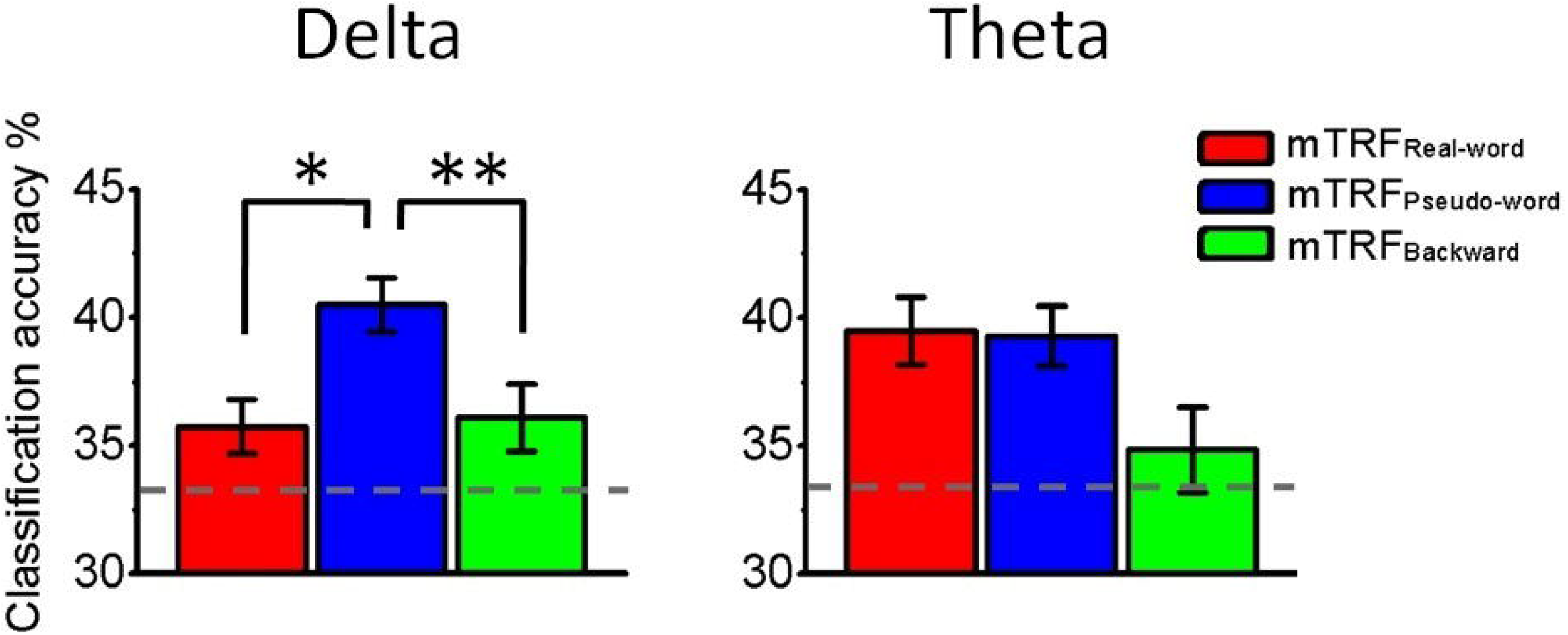
Accuracies of mTRF classification among all stimulus types (Scenario_1). The mTRF of each given stimulus types were tested by trials from all three stimulus types. Accuracies were based on *PredPowers* averaged over the centro-frontal electrodes. Statistical significance were Bonferroni corrected by the factor of 3. Dashed lines indicate the chance level (33.33%). Errors bars denote SEMs. * = significance at *p* < 0.05; ** = significance at *p* < 0.01.

**Fig. 7.**
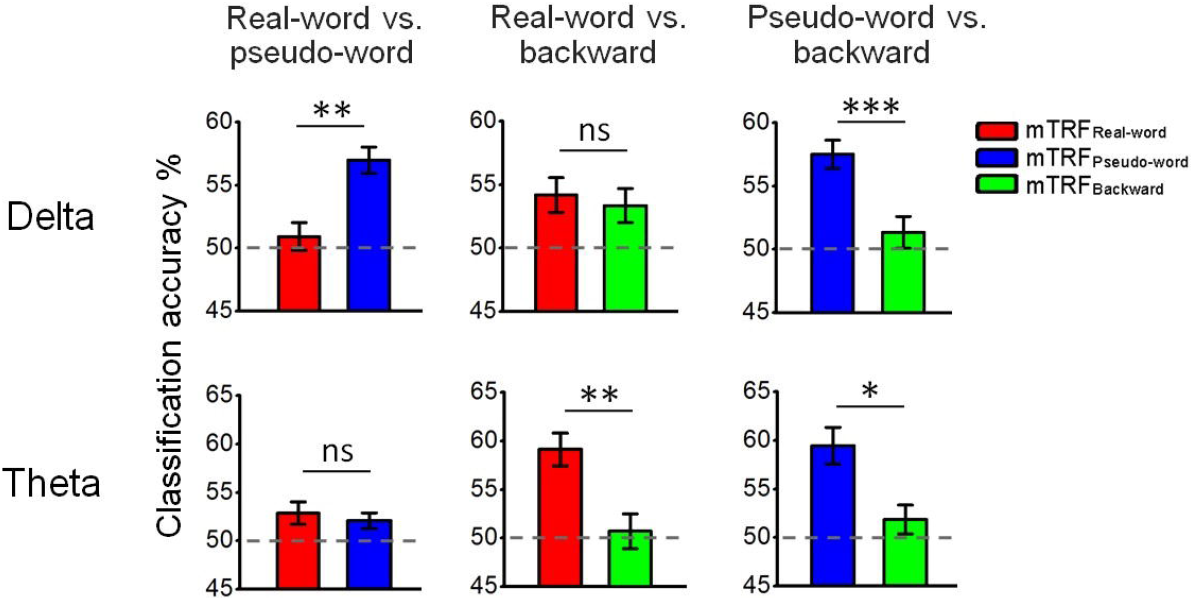
Accuracies of mTRF classification between any two stimulus types (Scenario_2). The mTRF of each given stimulus type was tested by trials from two stimulus types (one from the same stimulus type, the other from a different stimulus type). The accuracies were based on *PredPowers* averaged over the centro-frontal electrodes. Statistical significance were Bonferroni corrected by the factor of 3. Dashed lines indicate the chance level (50%). Errors bars denote SEMs. * = significance at *p* < 0.05; ** = significance at *p* < 0.01; *** = significance at *p* < 0.001.

**Fig. 6** shows the accuracies of classification among all stimulus types (Scenario_1). The mTRF of each given stimulus type was tested by trials from all three stimulus types. Repeated Measures ANOVAs were conducted with the factor of Stimulus Type. Main effects of Stimulus Type were found at both delta and theta bands (see **Table 2**, mTRF classification in Scenario_1). Post-hoc tests found that, at the delta band, accuracies were significantly higher for pseudo-word than for real-word and backward utterances, while no significant difference was found between those for real-word and backward utterances (see **Fig. 6** and **Table 3**, delta-band mTRF classification in Scenario_1). At the theta band, no significant differences of accuracies were found between any two stimulus types (see **Fig. 6** and **Table 3**, theta-band mTRF classification in Scenario_1).

**Fig. 7** shows the accuracies of classification between any two stimulus types (Scenario_2). The mTRF of each given stimulus type was tested by trials from two stimulus types, one from the same stimulus type and the other from a different stimulus type. We tested three pairs of comparisons: (1) real-word vs. pseudo-word (mTRF_Real-word_ and mTRF_Pseudo-word_ tested by trials from both real-word and pseudo-word utterances); (2) real-word vs. pseudo-word (mTRF_Real-word_ and mTRF_Backward_ tested by trials from both real-word and backward utterances); (3) pseudo-word vs. backward (mTRF_Pseudo-word_ and mTRF_Backward_ tested by trials from both pseudo-word and backward utterances). At the delta band, the results resembled those in Scenario_1, where accuracies were significantly higher for pseudo-word than for real-word and backward utterances, and there was no significant difference between real-word and backward utterances (see **Fig. 7** and **Table 3**, delta-band mTRF classification in Scenario_2). At the theta band, accuracies were significantly higher for real-word and pseudo-word than backward utterances, and no significant difference was found between real-word and pseudo-word utterances (see **Fig. 7** and **Table 3**, theta-band mTRF classification in Scenario_2).

### 3.5 Temporal properties of mTRFs

**Fig. 8** shows the spectrotemporal representations of mTRFs for different stimulus types averaged over the centro-frontal electrodes. Delta mTRFs (upper panels) showed temporal fluctuations that persisted across the entire 300-ms range for all stimulus types, while theta mTRFs (lower panels) showed N1-P1-N2-like complexes within the first 150 ms before the weighting reached a relatively low and stable level. **Fig. 9** shows the absolute mTRF weighting averaged across the 16 spectral bands. To examine how the degrees of neural entrainment vary across time lags for different stimulus types, Repeated Measures ANOVAs were conducted for the absolute weighting with the factors of Time (‘early’ (20 ~ 160 ms) vs. ‘later’ (160 ~ 300 ms)) and Stimulus Type. For delta-band absolute weighting, no significant main effects of Time or Stimulus Type, or [Time × Stimulus Type] interaction were found (see **Table 4**, delta-band). For theta-band absolute weighting, there was a significant main effect of Time, but no significant main effect of Stimulus Type or [Time × Stimulus Type] interaction (see **Table 4**, theta-band). Theta-band absolute weighting was significantly greater at the ‘early’ than at the ‘later’ time lags (see **Fig. 9**). The statistical results are thus consistent with the features of mTRF series shown in **Fig. 8**.

**Fig. 8.**
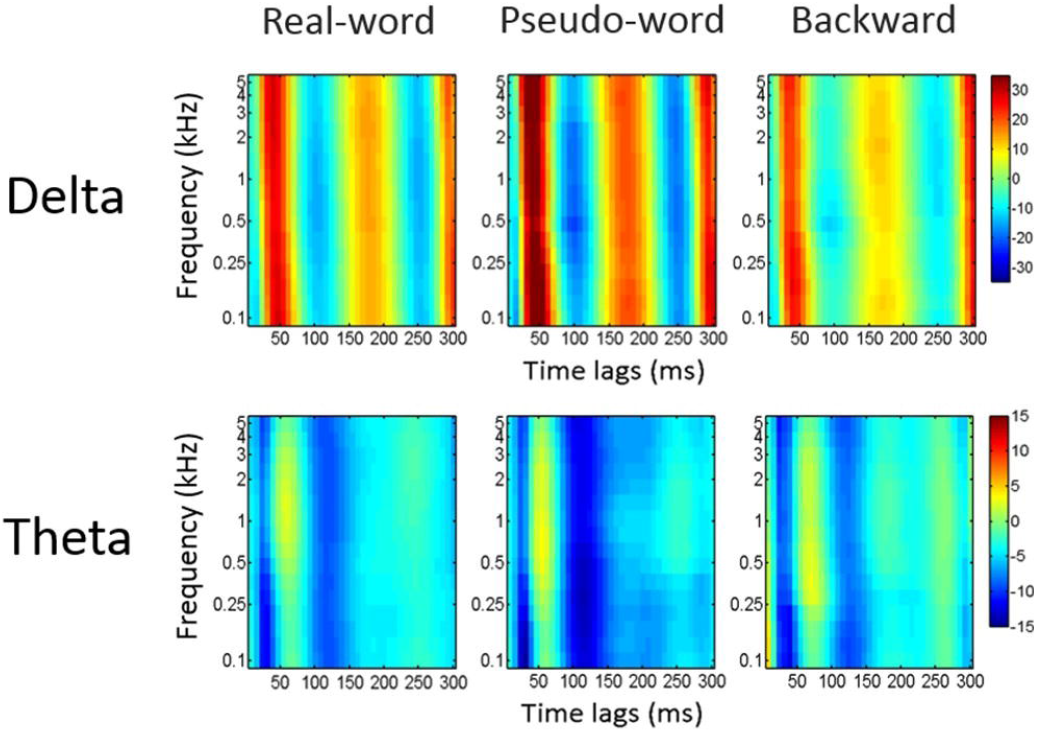
Spectrotemoral representations of mTRFs averaged over the centro-frontal electrodes. Note that frequencies are in logarithmic scale divided into 16 spectral bands (see *Methods*).

**Fig. 9.**
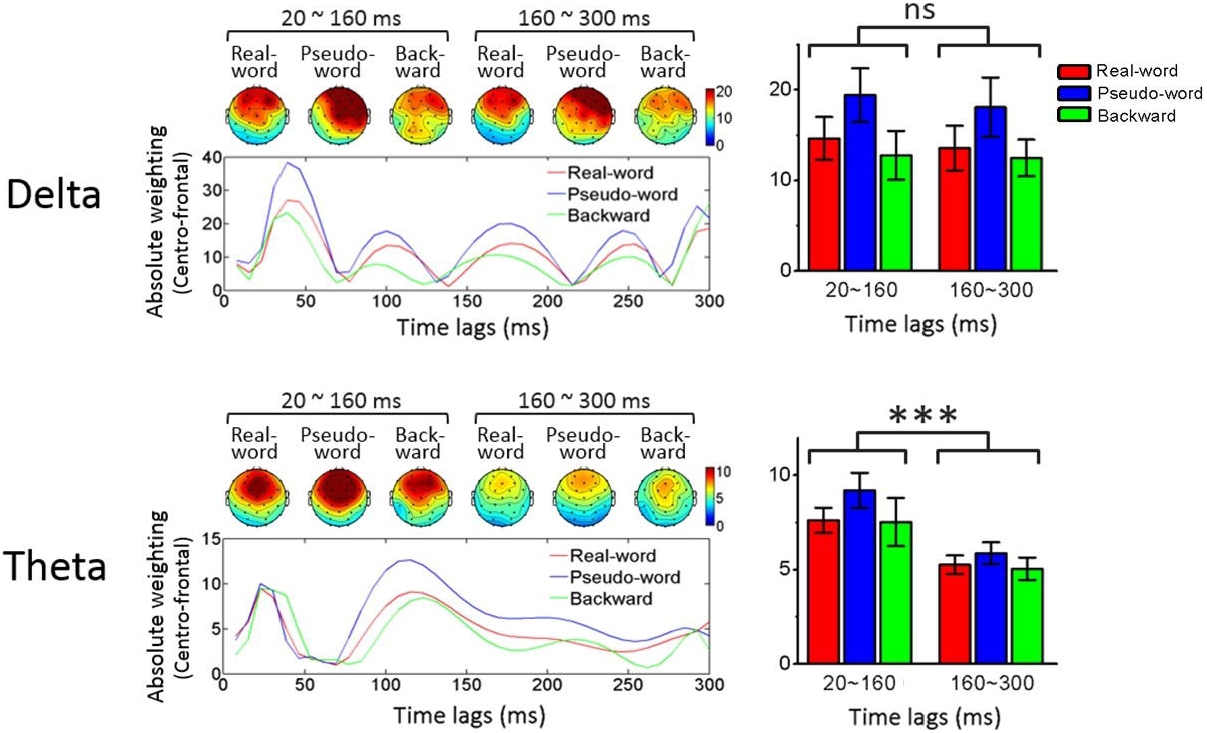
Absolute weighting of mTRFs. Left panels show the time series of absolute weighting averaged over the 16 spectral bands and the centro-frontal electrodes (the line graphs) and the corresponding scalp topographies of the ‘early’ (20 ~ 160 ms) and ‘later’ (160 ~ 300 ms) time lags for different stimulus types. Right panels show the comparisons of the absolute weighting between time lags (‘early’ vs. ‘later’). Errors bars denote SEMs. *** = significance at *p* < 0.001; ns = not significant.

**Table 4.** Statistical results of Repeated Measures ANOVAs for absolute weighting of mTRFs (averaged over the centro-frontal electrodes) across Time (‘early’ vs. ‘later’) and stimulus types. *Df, F, p* and *η_p_^2^* refer to degrees of freedom, F-values, *p*-values and partial eta-squared, respectively. The statistics were Greenhouse-Geisser corrected. Numbers are all rounded to three decimal places, unless they are < 0.001. Significant *p*-values are indicated in bold. *** = significance at *p* < 0.001.

| Dependent variables | Band | Factors | $df1$ | $df2$ | $F$ | $p$ | $\eta_p^2$ |
| --- | --- | --- | --- | --- | --- | --- | --- |
| Absolute weighting of mTRFs | Delta | Time | 1 | 19 | 0.811 | 0.379 | 0.041 |
|  |  | Stimulus Type | 1.712 | 32.531 | 2.182 | 0.135 | 0.103 |
| | | Time × Stimulus Type | 1.573 | 29.893 | 0.131 | 0.829 | 0.007 |
|  | Theta | Time | 1 | 19 | 43.675 | **$10^{-5}$\*\*\*** | 0.697 |
|  |  | Stimulus Type | 1.899 | 36.076 | 1.151 | 0.326 | 0.057 |
| | | Time × Stimulus Type | 1.768 | 33.583 | 0.464 | 0.609 | 0.024 |

### 3.6 Result summary

The results are summarized in **Table 5**. We first showed that *PredPowers* were statistically greater when using mTRFs compared to univariate TRFs, consistent with the pervious finding (Di Liberto et al., 2015). We then confirmed that *PredPowers* based on mTRFs were above random level and tested the effectiveness of EEG encoding congruent stimulus envelope information. Congruent *PredPowers* were statistically greater than surrogate *PredPowers* for pseudo-word utterances at the delta band and for speech (real-word and pseudo-word) utterances at the theta band.

**Table 5.**
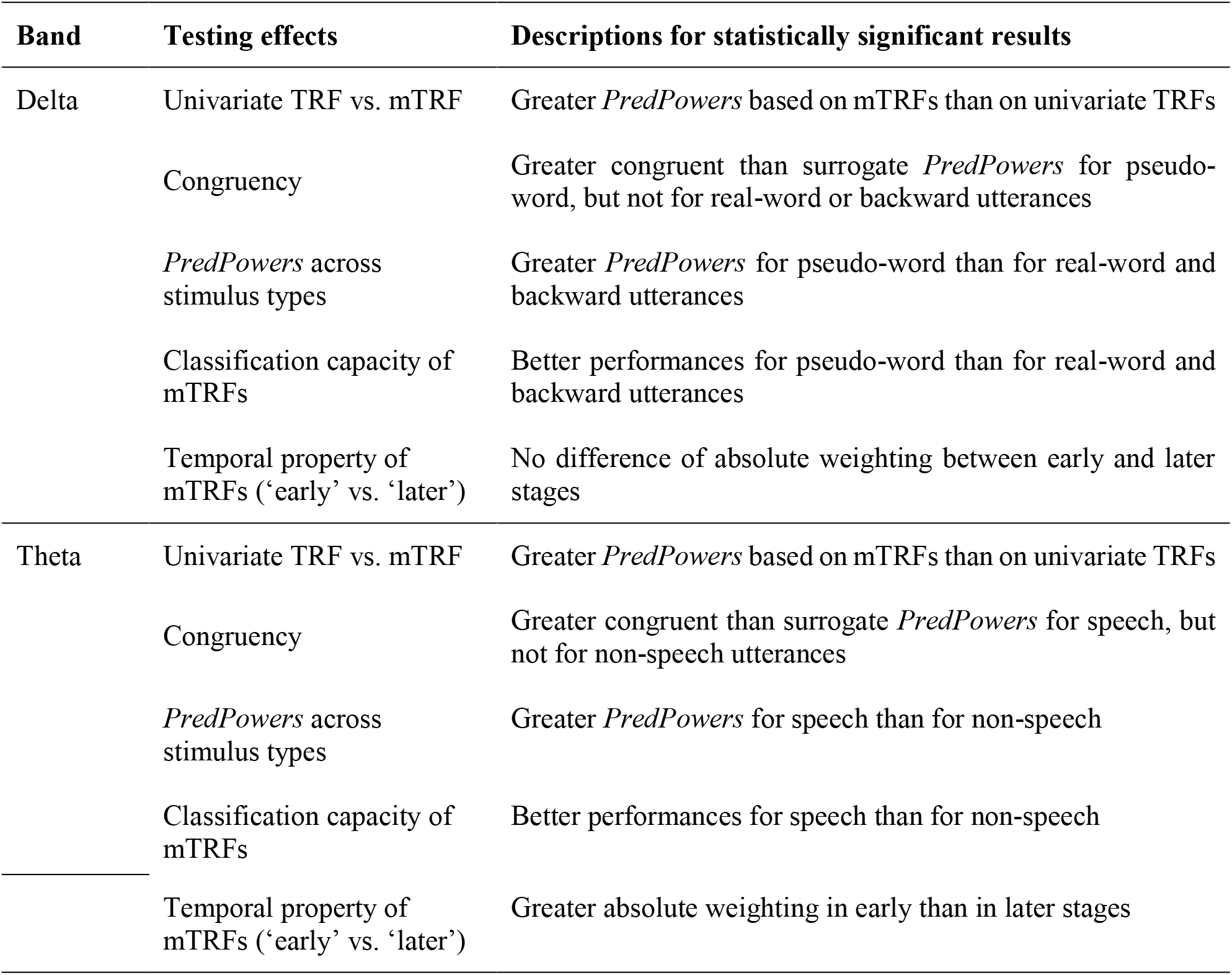
Brief summary of the results. ‘Speech’ refers to both real-word and pseudo-word utterances, while ‘non-speech’ refers to backward utterances.

*PredPowers* and classification capacity of mTRFs were then compared across stimulus types. The results showed a consistent pattern that delta- and theta-band entrainment take differential roles at different linguistic hierarchical levels. Specifically, delta-band *PredPower* was significantly greater for pseudo-word than for real-word and backward utterances, while theta-band *PredPower* was significantly greater for speech than for non-speech (backward) utterances. Correspondingly, delta-band mTRF had significantly better performances for pseudo-word than for real-word and backward utterances, whilst theta-band mTRF had significantly better performances for speech than for non-speech utterances.

We finally examined the temporal properties of the mTRF series, showing that the absolute weighting of mTRF at the theta, but not delta band, was significantly greater at early time lags (20 ~ 160 ms) than at later time lags (160 ~ 300 ms). This indicated that delta-band entrainment is likely to maintain across neural processing stages up to 300 ms, while theta-band entrainment mainly occurs at early stages of neural processing (< 160 ms).

## 4. Discussions

### 4.1 Superiority of mTRFs to univariate TRFs

We used a multivariate linear transformation algorithm that quantifies the neural entrainment of speech envelopes in EEGs (Di Liberto et al., 2015; Crosse et al., 2016). We showed that both delta- and theta-band *PredPowers* were significantly greater based on the mTRF than on the univariate TRF model. This is consistent with Di Liberto et al. (2015) which showed superiority of mTRF to univariate TRF when studying the low-frequency (1 ~ 15 Hz) neural entrainment of acoustic envelopes during speech perception. This indicates that low-frequency (both delta and theta) neural entrainment is achieved in a way that the brain encodes envelopes at multiple narrowbands at the cochlear output, rather than encodes the single broadband envelopes. While most previous MEG/EEG studies investigated the role of neural entrainment to single broadband acoustic envelopes for speech intelligibility (e.g., Peelle et al., 2013; Doelling et al., 2014; Vander Ghinst et al., 2016; Molinaro and Lizarazu, 2018; Vanthornhout et al., 2018), we suggest that the mTRF model provides a more appropriate approach of quantifying neural entrainment of acoustic envelopes during speech perception.

Note that, however, despite the superiority, a potential drawback of the mTRF model is that the linear mapping between envelopes and EEGs could be insensitive to the characteristics of response nonlinearities in audition (e.g., Christianson et al., 2008; Ahrens et al., 2008; Sadagopan and Wang, 2009). That being said, linear mapping is still a good approximation, as more advanced non-linear approaches in MEG/EEGs may yield greater computational complications and only marginal and negligible improvements (see detailed discussions by Crosse et al., 2016).

### 4.2 Distinctions between delta- and theta-band neural entrainment at different linguistic hierarchical levels

#### 4.2.1 Delta- and theta-band entrainment across stimulus types

Envelope modulations are critical acoustic cues for speech understanding (Drullman et al., 1994; Shannon et al., 1995; Arai et al., 1999; Swaminathan and Heinz, 2012). Previous MEG and EEG studies have shown that low-frequency neural entrainment of speech envelopes is associated with speech intelligibility (Peelle et al., 2013; Doelling et al., 2014; Vanthornhout et al., 2018). Recent tACS studies further showed that manipulating the degree of neural entrainment to envelopes can alter speech intelligibility, arguing the causal effect of the entrainment during speech perception (Zoefel et al., 2017; Riecke et al., 2018; Wilsch et al., 2018). Despite these findings, it is not clear, however, what is the role of neural entrainment at different linguistic hierarchical levels. Understanding speech should include processes of recognizing both phonological and semantic information (Nahum et al., 2008). Simply seen from the relationship between neural entrainment and speech intelligibility, how the entrainment subserves phonological and semantic processing during speech perception is still obscure. The present study therefore tested the EEG entrainment to speech envelopes in response to stimuli of real-word, pseudo-word and backward utterances that were expected to successfully dissociate the phonological and semantic processing (Binder et al., 2000; Londei et al., 2010; Saur et al., 2010; Mai et al., 2016). We found that delta-band neural entrainment (*PredPower*) was significantly greater for pseudo-word than for real-word and backward utterances. Theta-band neural entrainment, on the other hand, was significantly greater for speech (real-word and pseudo-word) than for non-speech (backward) utterances, but did not differ statistically between real-word and pseudo-word.

The result thus indicates the different roles that delta- and theta-band entrainment take during phonological and semantic processing. Greater theta-band entrainment for speech than for non-speech indicate its role in speech-specific processing, even though it can also occur in non-speech stimuli (theta-band *PredPower* for backward utterances was also above random level; see **Fig. 4**). The speech-specificity was likely to be associated with phonological, but not higher-level (semantic) processing, as it did not differ between real-word and pseudo-word utterances. This could indicate the neural tracking of syllabic- and sub-syllabic pattern of the speech signals during phonological processing. The delta-band entrainment, on the other hand, showed a distinct pattern, where the speech-specific properties were exhibited for pseudo-word utterances but not for real-word utterances. Plausibly, this may be explained by interactions between phonological and semantic processing during tracking of supra-syllabic rhythms, i.e., richer semantic information in the real-word utterances assisted in recognition of phonological contents, thereby reducing the demands of phonological processing indexed by the delta-band entrainment (Mai et al., 2016). From the perspective of pseudo-word utterances, delta-band entrainment was stronger possibly because of greater listening effort for phonological recognition due to lack of assistance from semantic information. This is in line with the behavioral studies showing the importance of delta-band envelopes for recognition of semantically meaningless syllables (Arai et al., 1996; 1999). It is also compatible with findings showing increased delta-band entrainment in some attention-demanding conditions, such as recognition of speech with reduced spectral resolution (Ding et al., 2014) or with increasingly noisy backgrounds (Vander Ghinst et al., 2016).

#### 4.2.2 Specificity of delta- and theta-band mTRFs for different stimulus types

*PredPowers* were compared between the ‘congruent’ and ‘surrogate’ conditions to test the specificity of mTRFs for different stimulus types. The congruency effect was found for pseudo-word but not for real-word or backward utterances at the delta band, and for speech but not for non-speech at the theta-band. Correspondingly, classification capacity of mTRFs were tested to see how congruent testing trials were accurately identified. Performances of delta-band mTRFs were better for pseudo-word than for real-word and backward utterances, while performances of theta-band mTRFs were better for speech than non-speech. Same patterns were shown in Scenario_1 and Scenario_2, although no statistical significance between any two stimulus types was found at the theta band in Scenario_1 (**Fig. 6** right panel). Lack of significance here may be because of the inability of theta-band mTRFs to distinguish testing trials between real-word and pseudo-word utterances, thereby decreasing the classification accuracies for both real-word and pseudo-word utterances when mTRFs were tested by trials from all stimulus types. These results are thus consistent with the findings which compared entrainment between stimulus types, showing the highest specificity of delta-band entrainment for pseudo-word utterances and higher specificity of theta-band entrainment for speech than for non-speech.

Our results echo the recent MEG study showing that delta- and theta-band entrainment play different roles during speech perception (Molinaro and Lizarazu, 2018). Despite this, however, the roles of delta- and theta-band entrainment found in the present study were different from those in Molinaro and Lizarazu (2018). Molinaro and Lizarazu (2018) compared neural entrainment between speech and non-speech (2-Hz and 7-Hz AM white-noise, or spectrally-rotated speech). They found that, delta-band entrainment was greater for speech than for non-speech in the right superior temporal and left inferior frontal regions, while theta-band entrainment did not differ between speech and non-speech. It was therefore argued that delta-entrainment involves higher-order computations for language processing, while theta-entrainment involves perceptual processing of auditory inputs (Molinaro and Lizarazu, 2018). In contrast, our current results showed that greater delta-band entrainment for speech than for non-speech occurs only when semantic information are deficient (pseudo-word), while theta-band entrainment is greater for speech than non-speech regardless of the semantic contents.

There could be several reasons for the distinctions between our results and the findings by Molinaro and Lizarazu (2018). First, the non-speech stimuli used in Molinaro and Lizarazu (2018) were AM white-noise and spectrally-rotated speech with the same RMS intensity as the speech stimuli. In this case, due to huge differences of spectral distributions between speech and non-speech, perceptual loudness across spectral bands was not controlled. It may worth pondering whether such uncontrolled factor could influence the differences of neural entrainment between speech and non-speech. Our present study, on the other hand, used backward utterances as non-speech stimuli which kept the long-term spectrum the same as speech, thereby controlling the perceptual loudness across spectral bands. Second, in our present study, we used Mandarin utterances with the syllable rate controlled at ~ 4 Hz for all trials (see *Methods*), while it is not clear whether the syllable rate of the Spanish utterances was relatively fixed or varied across trials in Molinaro and Lizarazu (2018). We argue that the effect of neural entrainment at frequencies in the neighbourhood of the syllable rate (i.e., theta band) could be enhanced as a consequence of fixing the syllable rate across stimuli. This may be a possible reason for a stronger effect of theta-band entrainment in our present study. Third, different methods of quantifying neural entrainment were used. Molinaro and Lizarazu (2018) measured cross-spectral density between MEG signals and the broadband speech envelopes. The present study used linear transformation algorithms that involve training and testing mTRFs that reflect the extent of mapping between EEGs and speech envelopes from multiple spectral bands (Di Liberto et al., 2015; Crosse et al., 2016). Future work would be needed to clarify whether results obtained from different methods are comparable and consistent.

### 4.3 Different temporal properties between delta- and theta-band mTRFs

TRF can be seen as a fitting filter, through which acoustic features project to corresponding neural signals (Ding and Simon, 2012). TRFs can thus reflect the characteristics of how speech envelope information were encoded in the brain. Distinctions of temporal properties between delta- and theta-band mTRFs were found. Delta-band mTRFs showed persistent temporal fluctuations up to at least 300 ms, while early N1-P1-N2-like complexes were shown in theta-band mTRFs followed by gradual attenuations. Statistically, the absolute weighting of theta-, but not delta-band, mTRFs were greater at early (20 ~ 160 ms) than at later time lags (160 ~ 300 ms). It is likely that delta-band entrainment occurs not merely at early sensory processing stages, but also sustains during higher-order neural processing that take place later in time. Theta-band entrainment, on the other hand, has statistically higher probability to occur at early processing stages. Such properties are in line with the current finding that delta-, but not theta-band entrainment was affected by higher-level linguistic (semantic) information. This is also compatible with Molinaro and Lizarazu (2018) which argued that theta-band entrainment mainly reflects perceptual processing and delta-band entrainment involves additional higher-order processing during speech perception. In addition, similar to *PredPowers*, topographies of mTRFs showed dominant distributions over centro-frontal regions (**Fig. 9**), consistent with the previous finding (Crosse et al., 2015). Although source localization was not conducted due to extremely limited spatial resolution (32 electrodes), the relevant neural processing stages may involve a temporal-frontal cortical network during speech processing (Park et al., 2015; Molinaro and Lizarazu, 2018).

### 4.4 Possible effects of the behavioral tasks

Neural entrainment was tested when participants performed a forced-choice sound-matching task (see *Methods*). Such task may force participants to focus their attention on lower-level linguistic processing like phonological recognition. Previous studies have shown that behavioral tasks at different levels could alter the neural oscillatory activities. For example, Shahin et al. (2009) showed that EEG powers at theta to gamma bands were different between tasks of gender voice detection and semantic discrimination when participants listened to auditory words. McNab et al. (2012) showed that MEG powers at beta and gamma bands were modulated by tasks of phonological and semantic recognition in response to visual words. It is not clear whether neural entrainment is modulated by different behavioral tasks, which may needs to be studied further in the future.

### 4.5 Summary

The present study investigated the distinctions between delta- and theta-band neural entrainment of speech envelopes at different linguistic hierarchical levels, using auditory stimuli that dissociated phonological and semantic contents. Neural entrainment was measured using the mTRF model that mapped speech envelopes at multiple spectral bands onto EEGs. We demonstrated that theta-band entrainment was modulated by phonological, but not semantic contents, indicating its role of tracking syllabic and sub-syllabic patterns for phonological processing. Delta-band entrainment, on the other hand, was greater with rich phonological but deficient semantic information (pseudo-word). This may reflect the mechanism of interactions between phonological and semantic processing during tracking of supra-syllabic rhythms, i.e., reduced demands for phonological recognition in real-word utterances, or greater listening effort in pseudo-word utterances. Furthermore, through analysing temporal properties of mTRFs, we demonstrated that, delta-band entrainment sustained across neural processing stages up to ~ 300 ms, while theta-band entrainment is more likely to occur at early stages (< 160 ms).

Taken together, we confirmed our hypothesis that delta- and theta-band entrainment take distinct roles at different linguistic hierarchical levels. We suggest the results could improve our understanding and new insights into the mechanisms of neural encoding of acoustic features during speech perception in general. Further studies may be needed to clarify how different quantification methods and experimental tasks modulate the effect of neural entrainment.

## Acknowledgements

This study was supported in part by grant 455911 of the Hong Kong RGC to Prof. William S.-Y. Wang at The Chinese University of Hong Kong. Guangting Mai acknowledges support from the UCL Graduate Cross-Disciplinary Training Scholarship based at Departments of Experimental Psychology and Medical Physics & Biomedical Engineering.

## Footnotes

1 Although in Mandarin, a morphological valid syllable could convey certain semantic information, concatenating syllables without forming valid dyllabic words disrupts the semantic validity (c.f., Xiao et al., 2005), as in the pseudo-word utterances in the present study. All participants reported that they considered pseudo-word utterances as semantically invalid.

## Abbreviations

EEG: electroencephalography
MEG: magnetoencephalography
TRF: temporal response function
mTRF: multivariate temporal response function
tACS: transcranial alternative current stimulation
SUS: semantically unpredictable sentences
AM: amplitude-modulated

